# Behavioral Signature of Trihexyphenidyl in The *Tor1a* (DYT1) Knockin Mouse Model of Dystonia

**DOI:** 10.1101/2025.06.17.659360

**Authors:** Ahmad Abdal Qader, Yuping Donsante, Jeffrey E. Markowitz, H. A. Jinnah, Chethan Pandarinath, Ellen J. Hess

## Abstract

Dystonia is a neurological disorder characterized by involuntary repetitive movements and abnormal postures. Animal models have played a pivotal role in studying the pathophysiology of dystonia. However, many genetic models, e.g. the *Tor1a*+*/*Δ*E* (DYT1) mouse, lack an overt motor phenotype, despite significant underlying neuronal abnormalities within the striatum and other motor control regions. Because the striatum is implicated in action sequencing, it is possible that the behavioral defect arises as a disruption in the frequency and temporal ordering of behaviors, rather than execution, which cannot be captured using traditional behavioral assays, thus limiting drug discovery efforts. To address this challenge, we used MoSeq, an unsupervised behavioral segmentation framework, to compare the continuous free behavior of control *Tor1a*^+*/*+^ mice and knockin *Tor1a*^+*/*Δ*E*^ mutant mice in response to the anti-dystonia drug trihexyphenidyl. Although minimal baseline differences in behavioral organization were detected, both genotypes exhibited robust and consistent shifts in behavioral space structure after treatment with trihexyphenidyl. Further, we demonstrate differences in the behavioral space structure of male vs female mice after trihexyphenidyl challenge. The distinct behavioral signatures evoked by trihexyphenidyl and biological sex, a known risk factor for dystonia, suggest that the analysis of the temporal structure of continuous free behavior provides a sensitive and novel approach to the discovery of therapeutics for the treatment of dystonia.

## 1 Introduction

Dystonia is a family of motor abnormalities characterized by involuntary muscle contractions causing sustained or transient twisting movements, abnormal postures, or repetitive motions. The etiology is heterogeneous, including genetic mutations and brain injuries, but most dystonias are idiopathic and are more common in women than in men [1, 2, 3]. This complex heterogeneity has led to the creation of various genetic, pharmacological, and lesion-induced animal models of dystonia [4, 5] which have, in turn, provided insight into the underlying neuronal dysfunction. Notably, *TOR1A* (DYT1) dystonia is often used as a model system for understanding both the defects underlying dystonia and the mechanism of action of trihexyphenidyl, a small molecule drug frequently used to treat symptoms of dystonia. *TOR1A* dystonia is caused by a gag deletion (ΔE) in the *TOR1A* gene. Although it is a dominantly inherited disorder, only ∼30% of carriers express dystonia while most do not manifest overt motor dysfunction. Genetically engineered mouse models of *TOR1A* dystonia exhibit striatal dysfunction including aberrant cholinergic tone, reduced striatal dopamine transmission, and defects in corticostriatal synaptic plasticity [6, 7, 8, 9, 10, 11, 12, 13, 14], which are corrected by trihexyphenidyl (THP). Despite the neuronal dysfunction, these mice lack an overt behavioral phenotype and exhibit only subtle motor signs when challenged [15, 6], suggesting that the models are akin to non-manifesting carriers.

Because the striatum plays a critical role in the temporal organization of ethologically-relevant action sequences [16, 17, 18, 19] changes in striatal function caused by mutations, drugs or insult may result in covert deviations from normal behavioral repertoires that are not easily detected using traditional assays, which typically focus on a single behavioral task. Given the known striatal dysfunction in mouse models of *TOR1A* dystonia, these mice may be able to execute spontaneous actions indistinguishably from control mice, however, the stochastic order with which they transition from one action to another or the frequency at which they express a certain action may differ.

Likewise, THP, which is known to mediate striatal function, may elicit a behavioral signature. Such changes in behavior are not easily resolved via traditional behavioral assays because they are specifically designed to assess the performance of action execution. Thus, a comprehensive assay that assesses the overall continuous free mouse behavior while taking into consideration its temporal structure may be better suited to detecting deviations from typical behavior.

To determine if the *Tor1a*^+*/*Δ*E*^ (Δgag) mutation or THP alters the structure of spontaneous free behavior, we used MoSeq [20], an unsupervised behavioral tracking and segmentation framework grounded in the ethological observation that behavior is composed of organized probabilistic sequences of behaviors. MoSeq uncovers the collective space of sub-second behavioral motifs, or syllables, that constitute continuous free behavior in mice, and summarizes the statistics of how mice traverse that space. Further, it was previously demonstrated that MoSeq has sufficient resolution to detect underlying striatal dysfunction based on deviations in the sequencing of mouse behavior [16]. Here, we present an application of the MoSeq segmentation pipeline to 3D recordings of freely behaving control *Tor1a*^+*/*+^ and mutant *Tor1a*^+*/*Δ*E*^ mice. While the behavioral space structure of *Tor1a*^+*/*Δ*E*^ mice was similar to *Tor1a*^+*/*+^ mice, the behavioral space structure differed by sex and was significantly altered after treatment with THP, providing a robust behavioral signature that may be useful for the identification of therapeutics for the treatment of dystonia.

## 2. Materials and Methods

**Mice:** Heterozygous knockin mice carrying the *Tor1a(*Δ*gag*) mutation [21], and control littermates inbred on C57BL/6J were bred at Emory University. Mice were housed with a 12-hour light/dark cycle. Food and water were provided *ad libitum*. Mice were genotyped using PCR (forward primer 5’-GCTATGGAAGCTCTAGTTGG-3’; reverse primer 5’-CAGCCAGGGCTAAACAGAG-3’). All experimental protocols were approved by the Institutional Animal Care and Use Committee at Emory University and followed guidelines set forth in the Guide for the Care and Use of Laboratory Animals.

### Experimental Design

Seventeen mice (8-10 weeks of age) were tested: 10 mutant (5 males and 5 females) and 7 control littermates (3 males and 4 females). Mice were tested using a cross-over design with one week between test sessions. On the day prior to recording, mice were habituated in the open field arena for 30 minutes. On the following day, each mouse was injected subcutaneously with either saline, or 20 mg/kg trihexyphenidyl hydrochloride [TCI America], a dose that normalizes extracellular dopamine in mutant *Tor1a*^+*/*Δ*E*^ mice [22]. Treatments were administered in pseudorandom order. Mice were then placed in the open field arena for 10 minutes before recording of the freely-moving behavior commenced (Figure 1A, top). Each recording session lasted 30-35 minutes and occurred between 9:30am and 3pm. Mice that received saline in the first test session received THP in the second session, and vice versa.

**Figure 1:**
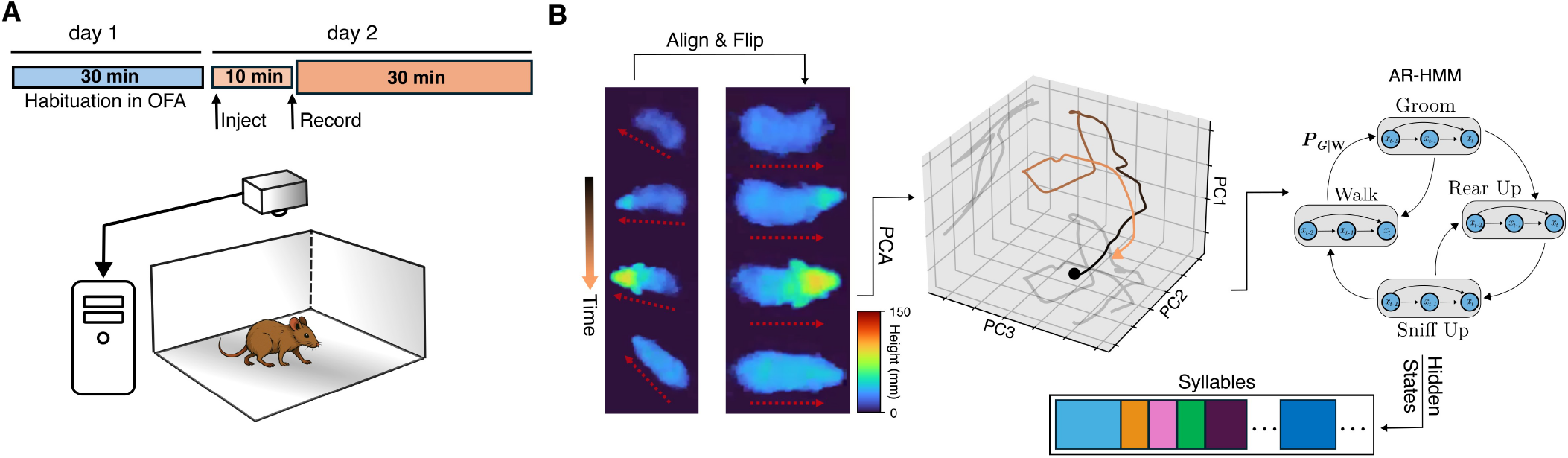
Illustration of the experimental apparatus and analysis pipeline. **(A)** Each mouse was habituated to the open field arena (bottom) for 30 minutes on the day prior to recording. On the following day, the mouse was injected, placed in the arena for 10 minutes before recording commenced. A camera directly above the arena recorded depth images and streamed them directly to disk. **(B)** Left: each row represents the extracted mouse from the recorded depth video from four time points throughout a rearing behavior both in the orientation as originally recorded (left column) and flipped to always face right (right column). The pseudocolor represents the height of the mouse. The dotted red arrows show the orientation of the mouse before and after the alignment and flipping procedure; middle: A sample trajectory in the top three PCs obtained by projecting the aligned, flipped mouse at every frame during the rearing behavior; right: the top 10 PCs are used as input data to a hierarchical autoregressive hidden Markov model (AR-HMM) whose hidden states represent the interpretable segmented behavioral syllables.

### Data Acquisition

Freely-moving mice were assessed in an open field that consisted of a 30×30×30 cm arena using a Helios 2+ time-of-flight depth sensor [Lucid Vision Labs, Canada] (Figure 1A, bottom) top-mounted 52 cm above the arena floor. Frames were acquired at 30Hz and each frame was 640×480 pixels, where each pixel’s value specifies the distance from the sensor. The acquisition clock was set via a hardware trigger using an Arduino UNO board. Data acquisition was managed using custom Python scripts running on a Linux machine to which uncompressed frames were directly streamed. Each recording session totaled approximately 30GB.

### Preprocessing & Data Preparation

All preprocessing was performed using custom Python scripts. For each recording, a rectangular region of interest that included only the arena boundaries was cropped to eliminate all other objects in the field of view. In each recording frame, all out-of-range pixel artifacts were replaced with ‘not-a-number’ values to be ignored and facilitate background isolation at later analysis stages. Pixel values were scaled to millimeters as 16-bit integers, and the resulting raw file was converted to a lossless .*avi* file for use in downstream steps.

### Analysis Pipeline

MoSeq is an unsupervised machine learning framework that segments free behavior and parses out discrete, sub-second stereotyped behavioral motifs, or syllables. It relies on the observation that continuous free behavior can be modeled as the organization of stereotyped action modules into sequences via a probabilistic selection mechanism [23, 20]. The version of MoSeq used here utilizes mouse 3D pose dynamics measured by a top-mounted depth camera. The full MoSeq analysis pipeline is detailed in [20, 24] and its publicly available code repository [25]. Briefly, the background of each depth recording was calculated as the median of a subset of filtered frames sampled at 500-frame intervals throughout the recording. The background was then subtracted to highlight the mouse. An expectation-maximization (EM) tracking model was used to identify the mouse in the presence of noise, such as dust particles close to the sensor, as described in [16]. In short, the EM model assumes that the mouse in a given depth frame can be well-approximated by a 3D Gaussian. In each frame, the Gaussian’s parameters were first estimated using the fit from the previous frame and then progressively optimized through iterative EM. The model’s outputs, which are likelihood-weighted pixels, were subsequently used to determine the mouse’s center and orientation via an ellipse fit. This process extracts the mouse from the surrounding arena and enables kinematics calculations such as body-averaged 3D speed and height at each frame. The speed corresponds to the motion of the center of the ellipse while the height is calculated as the average height across all the pixels within the ellipse.

Next, in each frame, an 80×80 pixel box centered around the mouse was cropped and aligned along the major axis of the ellipse, i.e., the spinal axis, such that the mouse was aligned horizontally (Figure 1B, left). A random forest model was then used to automate the detection of the mouse orientation (nose facing left or right), and the cropped box was flipped accordingly to ensure that the mouse was always facing right by convention. Samples from the processed recordings were reviewed by a human observer to ensure there were no artifacts and that flips in orientation were minimal. This alignment process ensured an approximate pixel-body part mapping, thus allowing for meaningful dimensionality reduction. All aligned cropped frames across all recordings were used to learn a low-dimensional projection using PCA, and each recording’s cropped aligned frames were then projected onto the top 10 principal components forming a 10-dimensional time series summarizing the 3D pose trajectory over the course of the recording (Figure 1B, center). The resulting PCA-projected pose summaries, which explained 85% of the variance, were used to fit a generative hierarchical autoregressive (AR) hidden Markov model whose hidden states represent the discrete identity of the behavioral syllable expressed, where each of these states is further represented via a separate AR process that describes the stereotyped continuous pose dynamics in PC space. Model training was performed according to the MoSeq published workflows and code repository, as noted above.

### Environmental Variables and Noise

Reflections were observed on two walls of the arena when mice reared on them. Therefore, preprocessing included an arena mask to minimize the reflections. To minimize noise in the depth images before projecting into the PC space, the right-aligned frames were smoothed in space using a 2D Gaussian filter with standard deviations (*σx, σy*) = (3.25, 2.0) pixels, and in time using a median filter of size 5 frames (i.e. 167ms).

### Quantitative Discovered Syllable Validation

To validate that the states identified by MoSeq are coherent, distinct, and appropriately describe the dataset, we used a cross-likelihood validation metric as described in [20]. Briefly, cross-likelihood estimates how well the AR parameters associated with each hidden state can describe observations assigned to every other state. Values near or above 1 would indicate that the AR parameters of a state describe the observations associated with another state, while negative values indicate failure of the AR parameters of one state at describing the observations associated with another.

### Statistical Testing

After establishing that speed and distance values are normally distributed (using quantile-quantile plots), t-tests and ANOVAs with treatment as the repeated measure were performed using GraphPad Prism (version 10.4.1). We calculated bigram-normalized transition matrices, which represent the probability of a syllable occurring given the identity of the previous syllable, as the frequency of the syllable pair divided by the total occurrences of the first syllable. To compare these bigram-normalized transition matrices across groups, we used the Jensen-Shannon Divergence (JSD) metric [26] and assessed significance using permutation testing. For each pair of experimental groups, we computed the pairwise JSD between all session combinations across groups. The resulting JSD values were then averaged to obtain a mean observed JSD, representing the similarity between the two groups. Next, session group labels were randomly shuffled 10,000 times, recalculating the mean permuted JSD for each iteration. The p-value was determined as the fraction of permutations in which the permuted JSD was greater than or equal to the observed JSD.

## 3 Results

### Aggregate kinematic summaries between genotypes

We first compared the gross kinematics, namely speed and height, between control and mutant mice. By computing a probability distribution over all speed and height values exhibited by each mouse, we built an archetypal behavioral profile for each experimental group. This aggregate-level behavioral profile served as a preliminary screen to examine differences in the overall movement patterns between experimental groups and across single sessions. We found that saline-treated control and mutant mice displayed similar distributions of speeds and heights (Figure 2: A & B, left column), suggesting that mutant mice exhibit an aggregate kinematic profile similar to control mice, in line with previous findings [6, 27]. Specifically, the total distance traveled in 30 minutes did not differ significantly between saline-treated control and mutant mice (57.74 *±* 4.10 and 49.62*±*3.71 m, respectively, t_15_ = 1.450, *p* = 0.168). Likewise, the average speed did not show significant differences between the saline-treated control and mutant mice (29.972.20 and 25.76*±*1.99 mm/sec, respectively, t_15_ = 1.398, *p* = 0.182). This approach provides a broad overview of behavioral trends, but it does not offer insights into the sequential organization of behavior. That is, the patterns of behavioral syllables that produce continuous free behavior may differ across groups that might otherwise share similar gross kinematics features.

**Figure 2:**
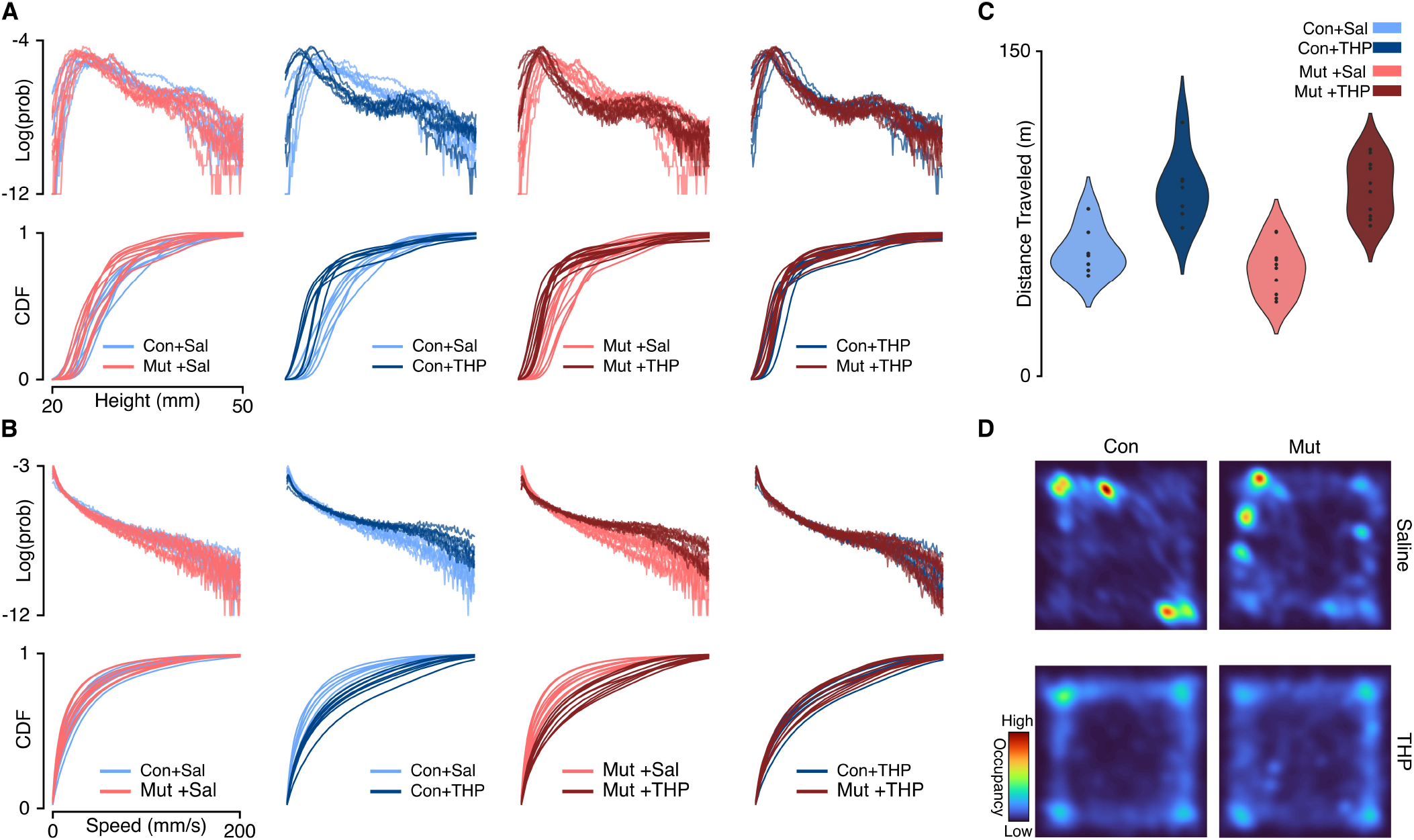
Single-session aggregate kinematics summaries compared across pairs of experimental groups. **(A)** Top: The log-transformed probability distribution of the exhibited height. Bottom: The cumulative distribution function of the exhibited height. Each column represents a comparison across a pair of experimental groups. Cool and warm palettes represent control (Con; n=7) and mutant (Mut; n=10) genotypes, respectively. Light and dark lines represent saline and THP treatments, respectively. **(B)** Same as **A** but representing the distributions over exhibited speeds. The log-transformation was used to facilitate visualizing the heavy-tailed distributions. **(C)** Total distance traveled in 30 minutes by each experimental group. **(D)** Representative single-session occupancy heatmaps of saline-treated (top) and THP-treated (bottom) mice; left column represents control mice (Con), right column represents mutant mice (Mut).

### Syllable expression frequency between genotypes

To resolve potential differences in the sequential and probabilistic structure of behavior between the two genotypes, we used MoSeq to uncover the sub-second behavioral syllables in our dataset to assess 1) how often control and mutant mice use each syllable and 2) the likelihood of transitioning from each syllable to every other syllable in each experimental group. MoSeq identified 44 coherent and stereotyped motifs, or syllables, that describe the behaviors in the dataset (Figure 3 and Supplementary Video 1). The syllables were visually inspected and further validated using a cross-likelihood estimate (Supplementary Figure 3) which showed effective separation of the syllables. At baseline, control and mutant mice exhibited similar syllable usage profiles (Figure 4A, top left).

**Figure 3:**
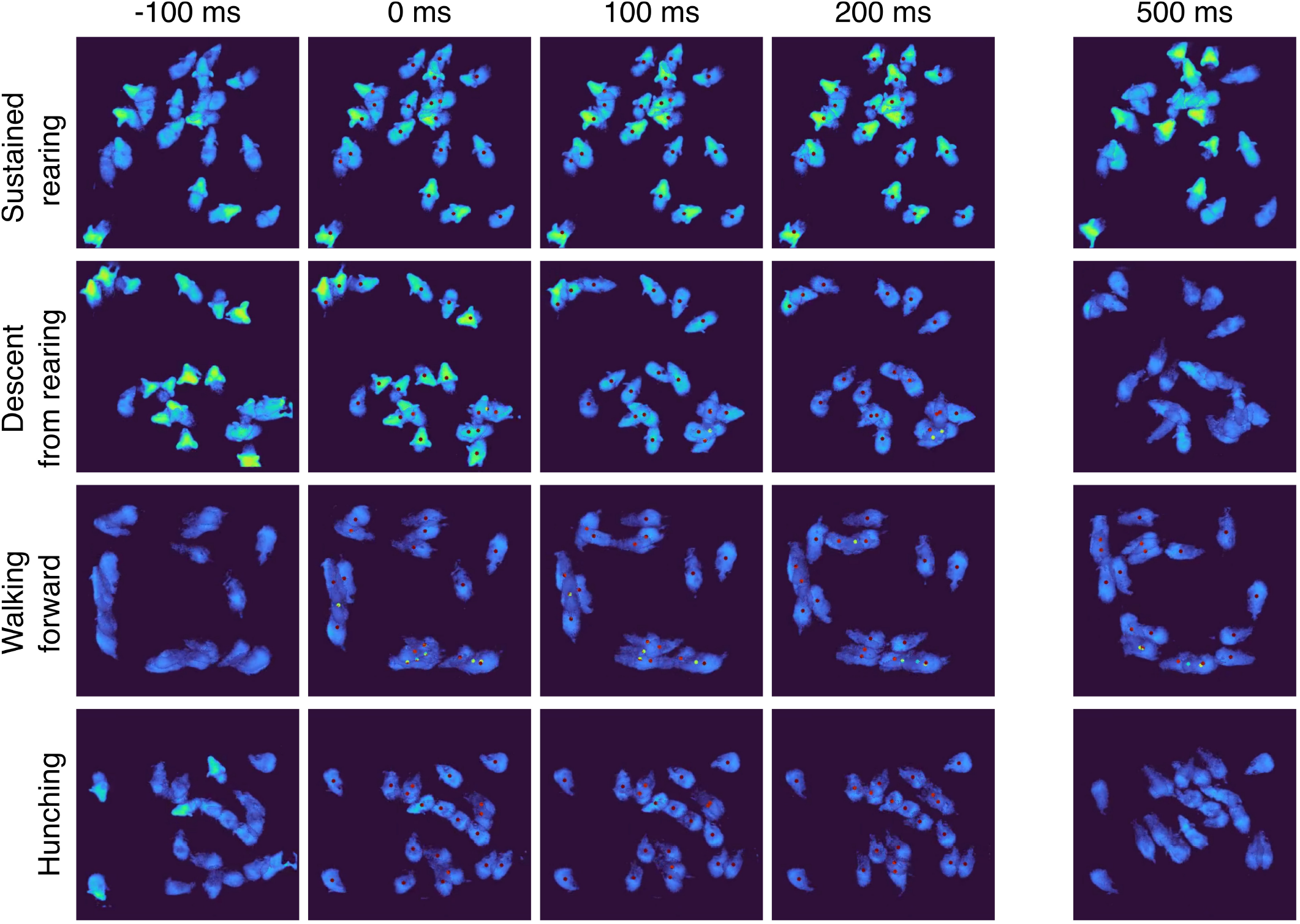
MoSeq identifies coherent stereotyped behavioral syllables. Montages of “crowd” movies depicting four behavioral modules, or syllables. A “crowd movie” is created by overlaying movies from individual mice exhibiting the same behavioral syllable. By superimposing multiple individual movies into one, the stereotyped nature of each behavioral syllable is highlighted. Here we show a 5-frame montage of “crowd” movies corresponding to four behavioral syllables. Each row represents a different syllable and columns represent progression in time at ~ 100ms increments with a discontinuity after 200ms. The active expression of the syllable is marked by a red dot placed over each “mouse instance”. Note that frames before the dot onset and after the dot offset can represent different behavioral syllables.

**Figure 4:**
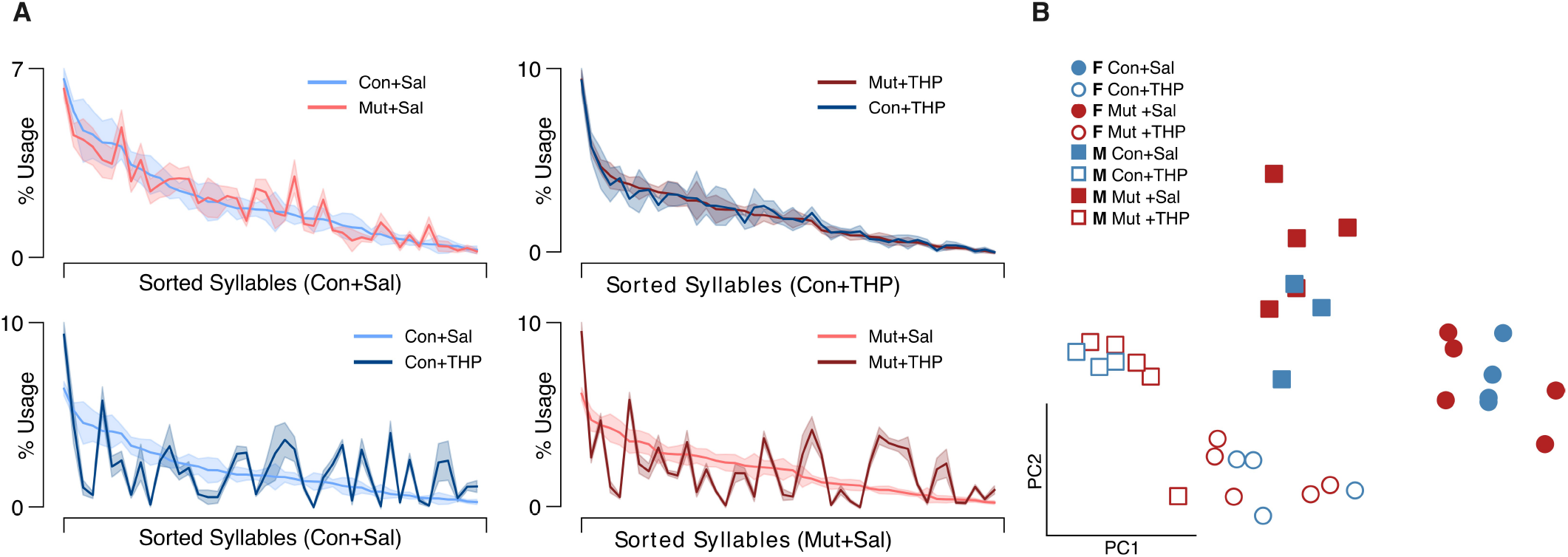
Syllable usage across genotype and treatment. **(A)** Average syllable usage profiles of experimental group pairs. The syllables are sorted in descending order based on usage by the control group indicated on the x-axis in each panel to highlight trend differences. Top row shows comparison across genotypes; bottom row shows comparison across treatments; shaded area represents the SEM. **(B)** Two-dimensional projection of single sessions’ empirically-obtained syllable usage vectors on the top two principal components; blue and red colors represent control (Con) and mutant (Mut) genotypes, respectively; squares and circles represent males and females, respectively.

### Syllable sequencing between genotypes

With access to the discrete behavioral syllables that describe our dataset and with a syllable identity assigned to each time point, it was possible to examine the sequential structure of mouse behavior across groups by computing a conditional probability distribution that quantifies the likelihood of transitioning from syllable A to any other syllable B. This distribution is captured by the bigram-normalized, empirically-observed transition matrix of a single session. To obtain a group-level representation, we averaged the observed transition matrices across mice within an experimental group. The resulting distribution was visualized as a transition graph where each node corresponds to a syllable and edges between nodes represent the transition probabilities (Figure 5A). To visually compare transition dynamics of a pair of groups, the corresponding transition graphs were subtracted to obtain a difference graph that highlights the differences in transition probabilities (Figure 5B). Visual inspection of the difference graphs suggested minimal differences between control and mutant mice. Further, quantitative assessment of the “distance” between transition probability distributions across groups, using the Jensen-Shannon divergence, showed no significant differences in transition dynamics between the two genotypes (Figure 5A).

**Figure 5:**
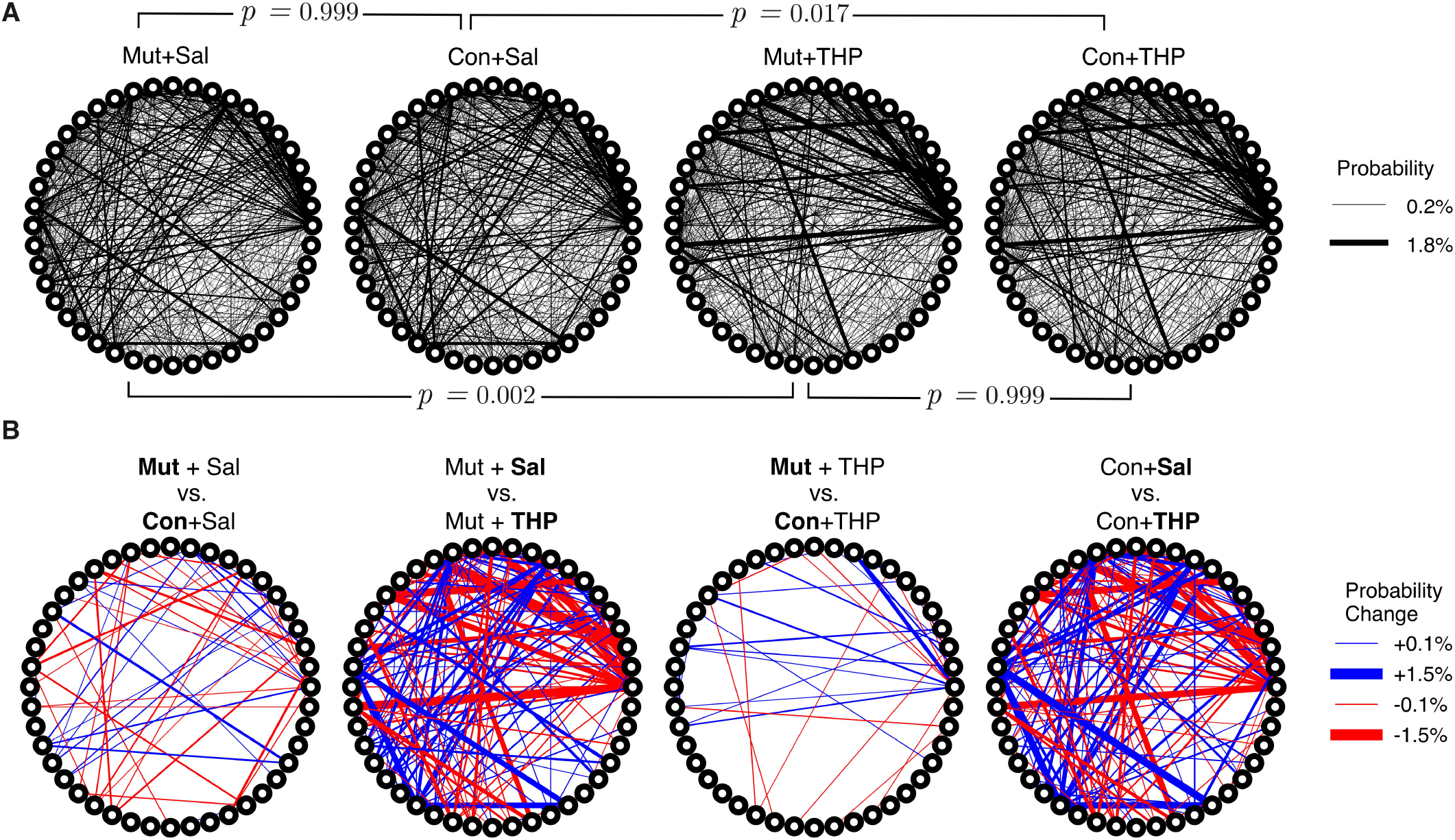
Differences in the temporal organization of syllable expression across experimental groups. **(A)** Transition graphs representing averaged transition dynamics in control (Con) and mutant (Mut) mice in response to saline or THP challenge. Each node on the circumference represents a syllable and edge thickness represents the corresponding bigram-normalized transition likelihood. Statistical testing was performed using a permutation test with a JSD statistic, brackets indicate pairs of groups compared. **(B)** Difference transition graphs obtained by subtracting a pair of group-averaged transition graphs to highlight differences; blue and red edges represent upregulation and downregulation of a particular transition, respectively.

**Figure 6:**
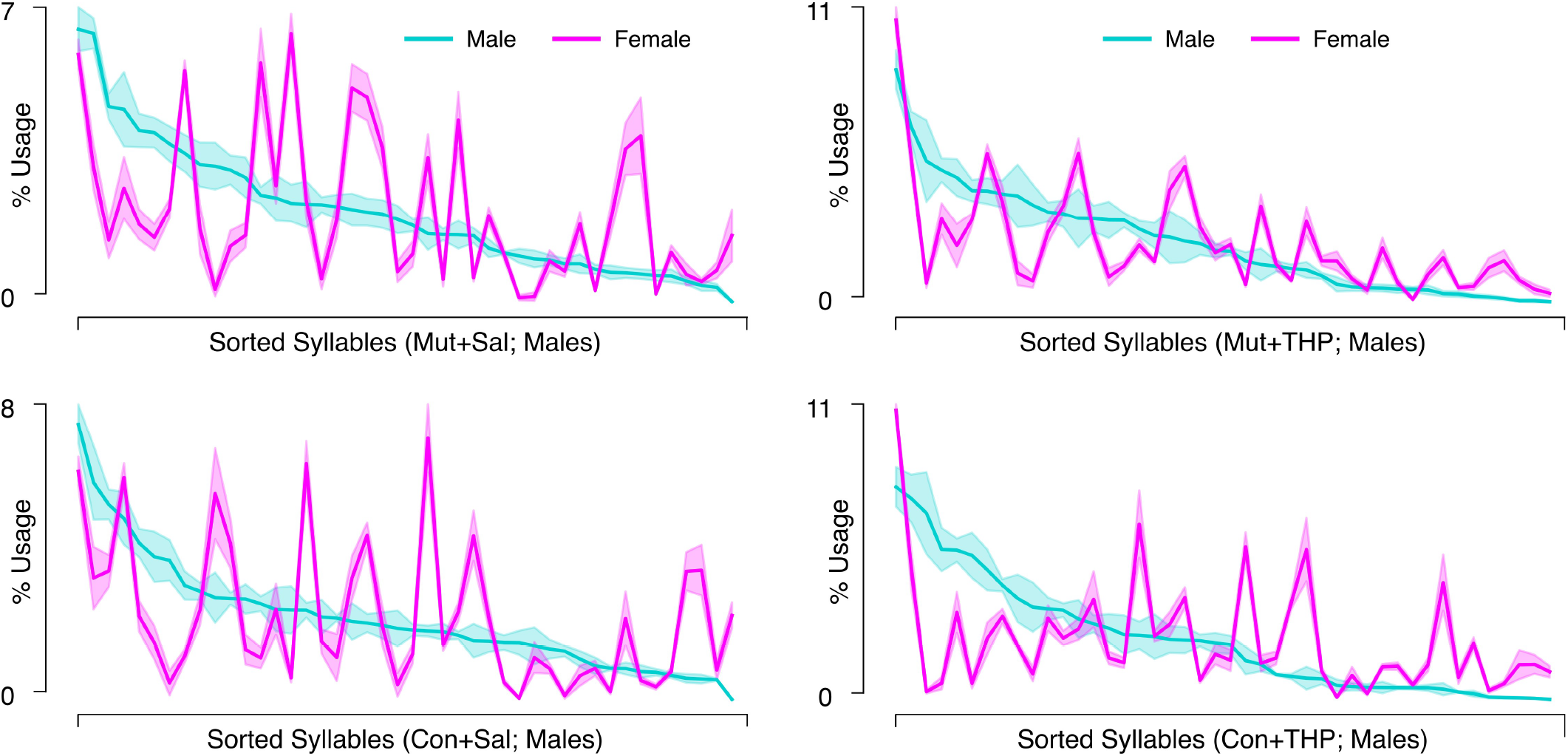
Syllable usage across sex within genotype-treatment combinations. Each panel corresponds to the average syllable usage profiles of a genotype-treatment combination specified in the x-axis label in control (Con) and mutant (Mut) mice. Syllables are sorted in descending order based on usage by male mice in that group. Cyan and magenta represent males and females, respectively. Shaded area represents the SEM.

### Behavioral space structure after THP challenge

After comparing the behavioral spaces of control and mutant mice, we then compared the effects of THP on behavioral composition. Both THP-treated control and mutant mice displayed vastly different speed and height distributions compared to saline treatment (Figure 2: A & B, middle columns), demonstrating that the effect of THP can be resolved at this statistical summary level. Consistent with prior studies [28, 29, 30], THP treatment induced a significant increase in locomotor activity compared to saline treatment (Figure 2 C) in both control and mutant (2-way repeated measures ANOVA, main effect of treatment, F_1,15_ = 159.4, *p <* 0.0001; main effect of genotype, F_1,15_ = 0.4774, *p* = 0.5001). Moreover, we found that THP augmented fast behaviors in both control and mutant mice, as reflected in the initial slow rise of the speed cumulative distribution functions (CDFs) (Figure 2B, middle columns) and the average speed (Supplementary Figure 1B) in control and mutant (2-way repeated measures ANOVA, main effect of treatment, F_1,15_ = 160.1, *p <* 0.0001; main effect of genotype, F_1,15_ = 0.4254, *p* = 0.5241). Additionally, THP-treated control and mutant mice displayed similar kinematics distributions (Figure 2: A & B, right column), suggesting that both genotypes exhibit comparable behavioral responses to THP at this aggregate kinematics level.

THP administration resulted in the reorganization of syllable usage profiles in control and mutant mice (Figure 4A, top right). This THP-induced restructuring was robust (Figure 4A, bottom row), reflecting a substantial reorganization of behavioral output in response to THP in both control and mutant mice. Despite these pronounced changes, MoSeq revealed minimal differences in syllable frequency between control and mutant mice in response to THP. Indeed, the transition graphs illustrate the THP-induced change in the behavior of both genotypes and also demonstrate the similarity of the behavior elicited by THP in both genotypes. Syllable usage profiles visualized in PC space (Figure 4B) similarly revealed distinct clusters between saline treatment and THP treatment while the genotypes overlapped within each cluster. Surprisingly, we also observed a separation between males and females (with the exception of one THP-treated male) in PC space (described below). To ensure that the treatment effect was not dominating the top PCs and diluting the genotype effect, we performed Linear Discriminant Analysis (LDA) with sex, treatment, and genotype separately to maximize class separability. We observed substantial overlap between the two genotypes across sex and treatments (Figure S2), but notably, saline-treated males showed the largest separability compared to other groups. Additionally, quantitative assessment of the transition probability distributions showed no significant differences in transition dynamics between the genotypes regardless of treatment (saline or THP), whereas substantial differences were observed between saline baseline and THP challenge (Figure 5A) in both genotypes. In contrast to the PC analysis, difference transition graphs showed subtle differences in transition dynamics between the two genotypes in the saline baseline condition, but less so after THP administration (Figure 5B).

### Sex differences

The separation between males and females in PC space suggested underlying sex differences. Therefore, we compared distance traveled and kinematics between sexes in all experimental conditions. Females covered significantly more distance than males and the speed distributions suggested that females exhibited faster speeds than males after both saline and THP treatments (Supplementary Figure 1 A & C). Further, the THP-induced increase in speed appeared more pronounced in females than in males, suggesting that the effect of THP is potentiated in females.

## 4 Discussion

We used a state-of-the-art method to segment continuous free behavior of control and mutant mice in an objective and unsupervised manner. Both genotypes displayed similar speed and height distributions and appeared to share a similar behavioral repertoire although subtle differences that did not reach significance were detected. While males and females exhibited unique behavioral landscapes, obvious differences between the genotypes within sex were not observed. In contrast, THP, an anticholinergic drug used to alleviate dystonia, significantly shifted the baseline behavioral landscape, suggesting a novel approach for the identification of antidystonic drugs.

The similarity in the temporal sequencing of syllables between mutant and control mice is remarkable given the deficit in dopamine release in the mutant mouse striatum [22, 8], which has been shown to play a critical role in probabilistic action sequencing [17, 16, 18, 19]. While striatal lesions in adult mice result in overt behavioral changes, the *Tor1a* gene defect affects neuronal dysfunction throughout development [31] which may trigger a compensatory mechanism that stabilizes motor function and sequencing despite the ~50% reduction in dopamine release in mutant mice. This mechanism could be a layer of homeostatic plasticity at the level of spiny projection neurons or a larger, network-level compensation involving the cortex.

Our approach was extremely sensitive for capturing the behavioral signature of the antidystonic drug THP, validating the sensitivity of the assay to behavioral change. In addition to a pronounced locomotion-enhancing effect, consistent with previous findings in rodents [28, 29, 30], we found that THP augmented fast behaviors and reorganized syllable usage. This robust behavioral response was observed in control and mutant mice, indicating that both genotypes share a common behavioral response to THP. One potential mechanism underlying this increase in locomotion activity and speed could be an increase in dopamine release in response to THP. This is supported by previous work demonstrating that THP increases dopamine release both in control and mutant knockin mice [14].

In addition to the effects of THP, there was a significant difference in the behavioral space between males and females. Biological sex is a known risk factor for most idiopathic focal dystonias including blepharospasm, oromandibular dystonia, cervical dystonia and spasmodic dysphonia [32, 33, 34, 35, 36, 37]. For inherited generalized dystonias, male-predominance obviously occurs in X-linked disorders, such as X-linked dystonia-parkinsonism. However, in the largest study published to date, male-bias was not observed in dystonias with autosomal inheritance, including *TOR1A* and *THAP1* dystonias [38]. In contrast, female bias occurs in some dystonias with autosomal inheritance including DOPA-responsive dystonia (DRD), which is caused by mutations in genes critical for dopamine synthesis, and dystonia caused by pathogenic variants of GNAL, which encodes the G-protein coupled to striatal D1 dopamine receptors [39, 38]. Notably, both of these female-predominant inherited dystonias are caused by pathogenic variants that disrupt dopamine neurotransmission. Despite these well-recognized sex differences, biological sex is rarely considered when developing therapeutics for dystonia, in part, because behavioral differences between the sexes are often subtle or undetectable. MoSeq provides a robust tool for assessing sex-biased therapeutics.

The version of MoSeq we used relies on a low-dimensional projection of aligned sequential depth images of mice as input to the autoregressive Hidden Markov Model (AR-HMM). Subsequently, variation in size of the mice in the dataset may have an impact on the behavioral segmentation performance of the trained model if these variations are considerable. One potential failure mode of the model in presence of such variations could be segmenting a stereotyped behavior into multiple disjoint syllables based on inherent size rather than the underlying pose dynamics. Since female mice are generally smaller in size than males, this might limit analyses regarding sex differences using syllable usage and transition statistics. However, methods to mitigate such limitations exist, such as performing a size normalization step to scale all mice into a common reference size before training the model, or using keypoint-MoSeq [40], which relies on trajectories of tracked keypoints as input instead of depth pose dynamics thus overcoming size-related limitations.

There are few drugs that are effective for the treatment of dystonia and there is currently no ‘gold standard’ therapeutic. THP is frequently used to treat dystonia and is the only small molecule drug proven effective in a double-blind placebo-controlled study in people living with dystonia [41]. However, the dose-limiting side-effects associated with this non-selective muscarinic acetylcholine receptor antagonist curtail its use, leaving patients with few options. New therapeutics have been slow to reach patients, in part, because preclinical testing in animal models generally involves labor-intensive *ex vivo* physiologic or behavioral assays that require highly specialized technical expertise. Here, we have demonstrated that THP elicits a clear and reproducible behavioral signature in freely moving mice using a streamlined assay. It is not yet clear if the THP-induced behavioral landscape is predictive of antidystonic drug effects in humans because the antidystonia pharmacopeia is so limited. However, identification of the fundamental behavioral signature of antidystonic drugs in the context of biological sex using ethological behavior as a predictor, as described here, has the potential to accelerate drug discovery in much the same way that the Porsolt swim test facilitated the discovery of antidepressant drugs.

## Author Contributions

Conceptualization: A.AQ and E.J.H; Investigation: A.AQ, Y.D; Coding and Formal Analysis: A.AQ, J.E.M, E.J.H; Visualization: A.AQ, H.A.J, E.J.H; Writing – Original Draft: A.AQ and E.J.H; Writing – Review & Editing: A.AQ, Y.D, J.E.M, H.A.J, C.P, E.J.H; Funding Acquisition: E.J.H

## References

1. Turcano P and Savica R. Chapter 17 - Sex differences in movement disorders. Handbook of Clinical Neurology. Ed. by Lanzenberger R, Kranz GS, and Savic I. Vol. 175. Sex Differences in Neurology and Psychiatry. Elsevier, 2020 Jan :275–82. doi: 10.1016/B978-0-444-64123-6.00019-9

2. Meoni S, Macerollo A, and Moro E. Sex differences in movement disorders. en. Nature Reviews Neurology 2020 Feb; 16:84–96. doi: 10.1038/s41582-019-0294-x

3. Rafee S, O’Riordan S, Reilly R, and Hutchinson M. We Must Talk about Sex and Focal Dystonia. en. Movement Disorders 2021; 36:604–8. doi: 10.1002/mds.28454

4. Wilson BK and Hess EJ. Animal models for dystonia. en. Movement Disorders 2013; 28:982–9. doi: 10.1002/mds.25526

5. Meringolo M, Tassone A, Imbriani P, Ponterio G, and Pisani A. Dystonia: Are animal models relevant in therapeutics? Revue Neurologique. INTERNATIONAL SFN /SOFMA MEETING 2018 2018 Nov; 174:608–14. doi: 10.1016/j.neurol.2018.07.003

6. Song CH, Fan X, Exeter CJ, Hess EJ, and Jinnah HA. Functional analysis of dopaminergic systems in a DYT1 knock-in mouse model of dystonia. Neurobiology of Disease 2012 Oct; 48:66–78. doi: 10.1016/j.nbd.2012.05.009

7. Balcioglu A, Kim MO, Sharma N, Cha JH, Breakefield XO, and Standaert DG. Dopamine release is impaired in a mouse model of DYT1 dystonia. en. Journal of Neurochemistry 2007; 102:783–8. doi: 10.1111/j.1471-4159.2007.04590.x

8. Page ME, Bao L, Andre P, Pelta-Heller J, Sluzas E, Gonzalez-Alegre P, Bogush A, Khan LE, Iacovitti L, Rice ME, and Ehrlich ME. Cell-autonomous alteration of dopaminergic transmission by wild type and mutant (?E) TorsinA in transgenic mice. Neurobiology of Disease 2010 Sep; 39:318–26. doi: 10.1016/j.nbd.2010.04.016

9. Martella G, Maltese M, Nisticò R, Schirinzi T, Madeo G, Sciamanna G, Ponterio G, Tassone A, Mandolesi G, Vanni V, Pignatelli M, Bonsi P, and Pisani A. Regional specificity of synaptic plasticity deficits in a knock-in mouse model of DYT1 dystonia. Neurobiology of Disease 2014 May; 65:124–32. doi: 10.1016/j.nbd.2014.01.016

10. Pisani A, Martella G, Tscherter A, Bonsi P, Sharma N, Bernardi G, and Standaert DG. Altered responses to dopaminergic D2 receptor activation and N-type calcium currents in striatal cholinergic interneurons in a mouse model of DYT1 dystonia. Neurobiology of Disease 2006 Nov; 24:318–25. doi: 10.1016/j.nbd.2006.07.006

11. Grundmann K, Glöckle N, Martella G, Sciamanna G, Hauser TK, Yu L, Castaneda S, Pichler B, Fehrenbacher B, Schaller M, Nuscher B, Haass C, Hettich J, Yue Z, Nguyen HP, Pisani A, Riess O, and Ott T. Generation of a novel rodent model for DYT1 dystonia. Neurobiology of Disease 2012 Jul; 47:61–74. doi: 10.1016/j.nbd.2012.03.024

12. Martella G, Tassone A, Sciamanna G, Platania P, Cuomo D, Viscomi MT, Bonsi P, Cacci E, Biagioni S, Usiello A, Bernardi G, Sharma N, Standaert DG, and Pisani A. Impairment of bidirectional synaptic plasticity in the striatum of a mouse model of DYT1 dystonia: role of endogenous acetylcholine. Brain 2009 Sep; 132:2336–49. doi: 10.1093/brain/awp194

13. Sciamanna G, Tassone A, Mandolesi G, Puglisi F, Ponterio G, Martella G, Madeo G, Bernardi G, Standaert DG, Bonsi P, and Pisani A. Cholinergic Dysfunction Alters Synaptic Integration between Thalamostriatal and Corticostriatal Inputs in DYT1 Dystonia. en. Journal of Neuroscience 2012 Aug; 32:11991–2004. doi: 10.1523/JNEUROSCI.0041-12.2012

14. Downs AM, Fan X, Kadakia RF, Donsante Y, Jinnah HA, and Hess EJ. Cell-intrinsic effects of TorsinA(?E) disrupt dopamine release in a mouse model of TOR1A dystonia. Neurobiology of Disease 2021 Jul; 155:105369. doi: 10.1016/j.nbd.2021.105369

15. Dang MT, Yokoi F, McNaught KSP, Jengelley TA, Jackson T, Li J, and Li Y. Generation and characterization of Dyt1 ?GAG knock-in mouse as a model for early-onset dystonia. Experimental Neurology 2005 Dec; 196:452–63. doi: 10.1016/j.expneurol.2005.08.025

16. Markowitz JE, Gillis WF, Beron CC, Neufeld SQ, Robertson K, Bhagat ND, Peterson RE, Peterson E, Hyun M, Linderman SW, Sabatini BL, and Datta SR. The Striatum Organizes 3D Behavior via Moment-to-Moment Action Selection. English. Cell 2018 Jun; 174:44–58.e17. doi: 10.1016/j.cell.2018.04.019

17. Aldridge JW and Berridge KC. Coding of Serial Order by Neostriatal Neurons: A “Natural Action” Approach to Movement Sequence. en. Journal of Neuroscience 1998 Apr; 18:2777–87. doi: 10.1523/JNEUROSCI.18-07-02777.1998

18. Minkowicz S, Mathews MA, Mou FH, Yoon H, Freda SN, Cui ES, Kennedy A, and Kozorovitskiy Y. Striatal ensemble activity in an innate naturalistic behavior. en. eLife 2023 Apr; 12. doi: 10.7554/eLife.87042.1

19. Jin X, Tecuapetla F, and Costa RM. Basal ganglia subcircuits distinctively encode the parsing and concatenation of action sequences. en. Nature Neuroscience 2014 Mar; 17:423–30. doi: 10.1038/nn.3632

20. Wiltschko AB, Johnson MJ, Iurilli G, Peterson RE, Katon JM, Pashkovski SL, Abraira VE, Adams RP, and Datta SR. Mapping Sub-Second Structure in Mouse Behavior. English. Neuron 2015 Dec; 88:1121–35. doi: 10.1016/j.neuron.2015.11.031

21. Goodchild RE, Kim CE, and Dauer WT. Loss of the Dystonia-Associated Protein TorsinA Selectively Disrupts the Neuronal Nuclear Envelope. English. Neuron 2005 Dec; 48:923–32. doi: 10.1016/j.neuron.2005.11.010

22. Downs AM, Fan X, Donsante C, Jinnah HA, and Hess EJ. Trihexyphenidyl rescues the deficit in dopamine neurotransmission in a mouse model of DYT1 dystonia. Neurobiology of Disease 2019 May; 125:115–22. doi: 10.1016/j.nbd.2019.01.012

23. Tinbergen N. The Study of Instinct. en. Clarendon Press, 1951

24. Wiltschko AB, Tsukahara T, Zeine A, Anyoha R, Gillis WF, Markowitz JE, Peterson RE, Katon J, Johnson MJ, and Datta SR. Revealing the structure of pharmacobehavioral space through motion sequencing. en. Nature Neuroscience 2020 Nov; 23:1433–43. doi: 10.1038/s41593-020-00706-3

25. Datta Lab. MoSeq2 App. Available from: https://github.com/dattalab/moseq2-app.Commita09afb9. GitHub, 2021

26. Lin J. Divergence measures based on the Shannon entropy. IEEE Transactions on Information Theory 1991 Jan; 37:145–51. doi: 10.1109/18.61115

27. Sharma N, Baxter MG, Petravicz J, Bragg DC, Schienda A, Standaert DG, and Breakefield XO. Impaired Motor Learning in Mice Expressing TorsinA with the DYT1 Dystonia Mutation. en. Journal of Neuroscience 2005 Jun; 25:5351–5. doi: 10.1523/JNEUROSCI.0855-05.2005

28. Shimosato K, Watanabe S, and Kitayama S. Differential effects of trihexyphenidyl on place preference conditioning and locomotor stimulant activity of cocaine and methamphetamine. en. Naunyn-Schmiedeberg’s Archives of Pharmacology 2001 Jul; 364:74–80. doi: 10.1007/s002100100433

29. Tanda G, Ebbs AL, Kopajtic TA, Elias LM, Campbell BL, Newman AH, and Katz JL. Effects of muscarinic M1 receptor blockade on cocaine-induced elevations of brain dopamine levels and locomotor behavior in rats. eng. The Journal of Pharmacology and Experimental Therapeutics 2007 Apr; 321:334–44. doi: 10.1124/jpet.106.118067

30. Sipos ML, Burchnell V, and Galbicka G. Dose-response curves and time-course effects of selected anticholinergics on locomotor activity in rats. en. Psychopharmacology 1999 Dec; 147:250–6. doi: 10.1007/s002130051164

31. Li J, Levin DS, Kim AJ, Pappas SS, and Dauer WT. TorsinA restoration in a mouse model identifies a critical therapeutic window for DYT1 dystonia. en. The Journal of Clinical Investigation 2021 Mar; 131. doi: 10.1172/JCI139606

32. Williams L, McGovern E, Kimmich O, Molloy A, Beiser I, Butler JS, Molloy F, Logan P, Healy DG, Lynch T, Walsh R, Cassidy L, Moriarty P, Moore H, McSwiney T, Walsh C, O’Riordan S, and Hutchinson M. Epidemiological, clinical and genetic aspects of adult onset isolated focal dystonia in Ireland. en. European Journal of Neurology 2017; 24:73–81. doi: 10.1111/ene.13133

33. Hintze JM, Ludlow CL, Bansberg SF, Adler CH, and Lott DG. Spasmodic Dysphonia: A Review. Part 1: Pathogenic Factors. EN. Otolaryngology–Head and Neck Surgery 2017 Oct; 157:551–7. doi: 10.1177/0194599817728521

34. Pandey S and Sharma S. Meige’s syndrome: History, epidemiology, clinical features, pathogenesis and treatment. Journal of the Neurological Sciences 2017 Jan; 372:162–70. doi: 10.1016/j.jns.2016.11.053

35. The Epidemiological Study of Dystonia in Europe (ESDE) Collaborative Group. A prevalence study of primary dystonia in eight European countries. en. Journal of Neurology 2000 Oct; 247:787–92. doi: 10.1007/s004150070094

36. Marras C, Van den Eeden SK, Fross RD, Benedict-Albers KS, Klingman J, Leimpeter AD, Nelson LM, Risch N, Karter AJ, Bernstein AL, and Tanner CM. Minimum incidence of primary cervical dystonia in a multiethnic health care population. Neurology 2007 Aug; 69:676–80. doi: 10.1212/01.wnl.0000267425.51598.c9

37. Kilic-Berkmen G, Scorr LM, McKay L, Thayani M, Donsante Y, Perlmutter JS, Norris SA, Wright L, Klein C, Feuerstein JS, Mahajan A, Wagle-Shukla A, Malaty I, LeDoux MS, Pirio-Richardson S, Pantelyat A, Moukheiber E, Frank S, Ondo W, Saunders-Pullman R, Lohmann K, Hess EJ, and Jinnah H. Sex Differences in Dystonia. en. Movement Disorders Clinical Practice 2024; 11:973–82. doi: 10.1002/mdc3.14059

38. Lange LM, Junker J, Loens S, Baumann H, Olschewski L, Schaake S, Madoev H, Petkovic S, Kuhnke N, Kasten M, Westenberger A, Domingo A, Marras C, König IR, Camargos S, Ozelius LJ, Klein C, and Lohmann K. Genotype–Phenotype Relations for Isolated Dystonia Genes: MDSGene Systematic Review. en. Movement Disorders 2021; 36:1086–103. doi: 10.1002/mds.28485

39. Wijemanne S and Jankovic J. Dopa-responsive dystonia—clinical and genetic heterogeneity. en. Nature Reviews Neurology 2015 Jul; 11:414–24. doi: 10.1038/nrneurol.2015.86

40. Weinreb C, Pearl JE, Lin S, Osman MAM, Zhang L, Annapragada S, Conlin E, Hoffmann R, Makowska S, Gillis WF, Jay M, Ye S, Mathis A, Mathis MW, Pereira T, Linderman SW, and Datta SR. Keypoint-MoSeq: parsing behavior by linking point tracking to pose dynamics. en. Nature Methods 2024 Jul; 21. Publisher: Nature Publishing Group:1329–39. doi: 10.1038/s41592-024-02318-2

41. Burke RE, Fahn S, and Marsden CD. Torsion dystonia. Neurology 1986 Feb; 36:160–0. doi: 10.1212/WNL.36.2.160

