## Supplementary figures and images for "Behavioral Signature of Trihexyphenidyl in The *Tor1a* (DYT1) Knockin Mouse Model of Dystonia"

### Supplemental Figure 1

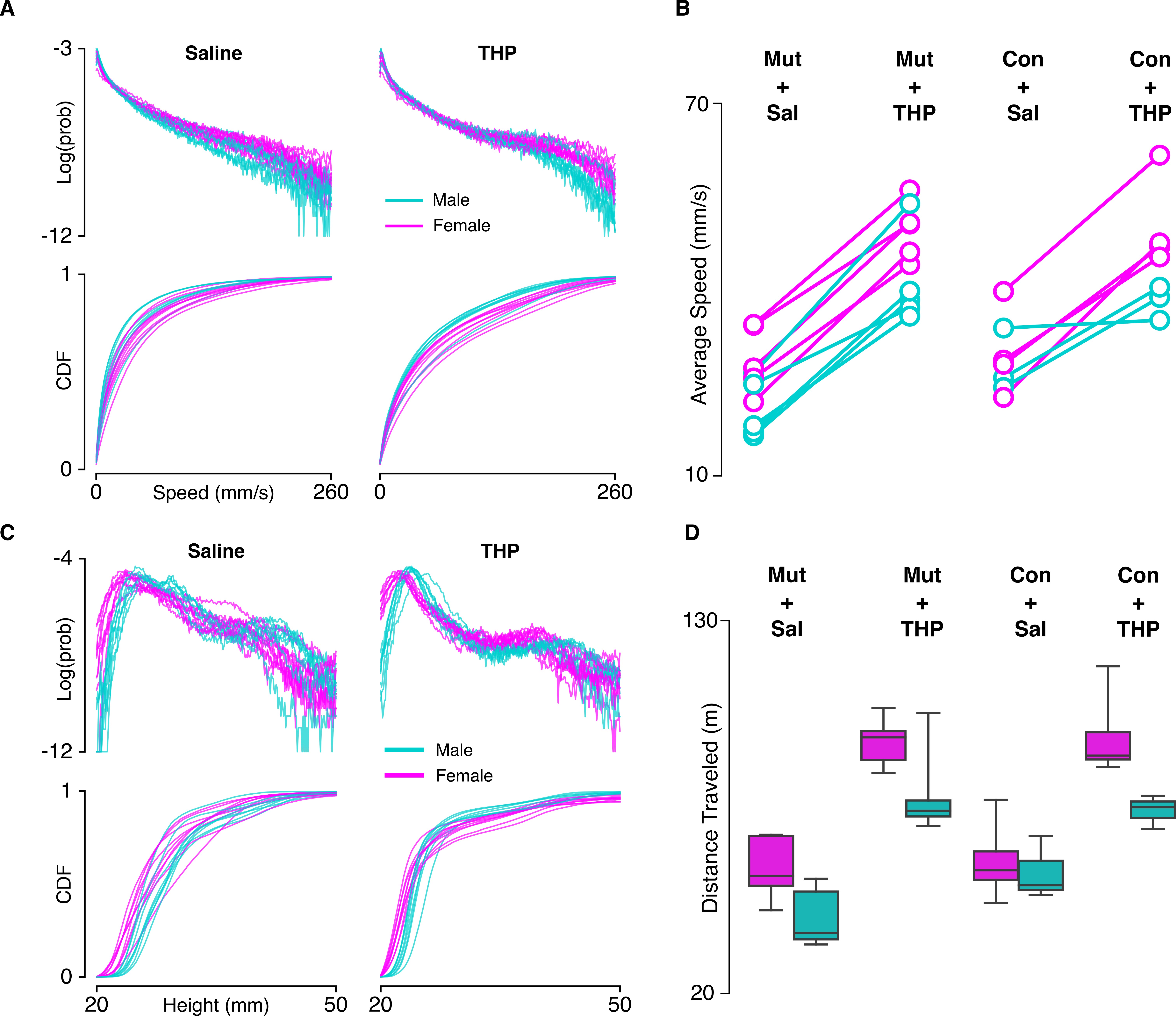

### Supplemental Figure 2

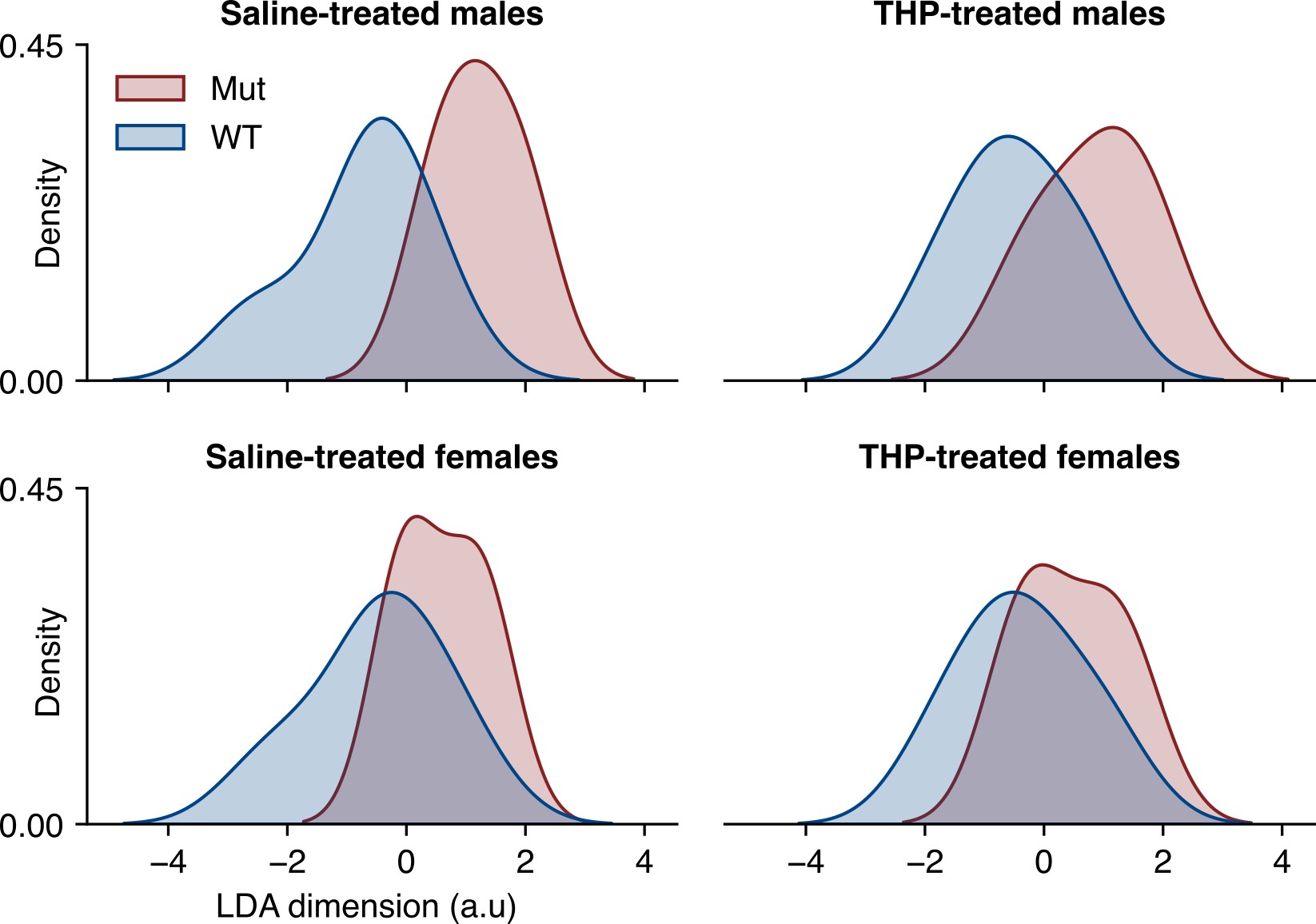

### Supplemental Figure 3

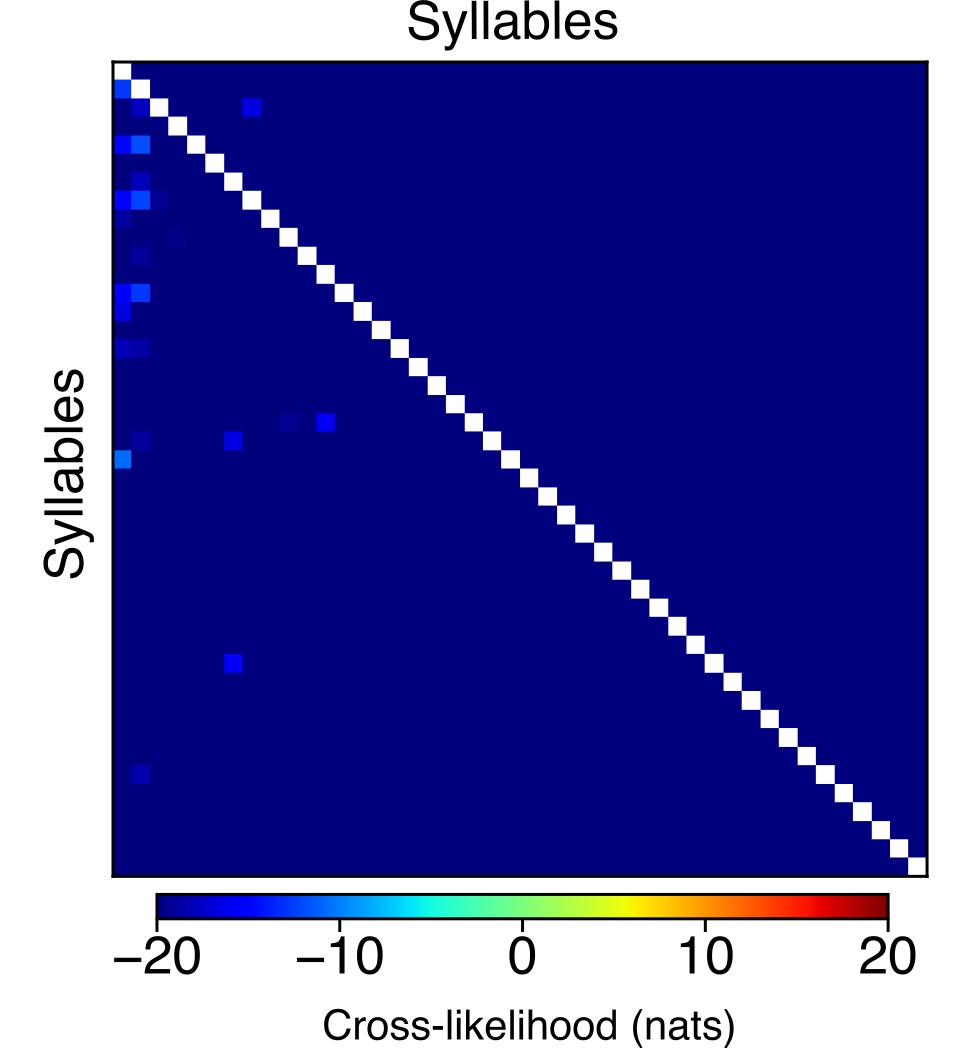
